# Does Low Dose Radiation Induced Adaptive Response Influence Initial DNA-DSB formation? Evidence from γH2AX foci Analysis in Human Lymphocytes

**DOI:** 10.64898/2026.05.19.726427

**Authors:** Sheeri Fatima, Aarti Notnani, Rajesh Kumar Chaurasia, Kapil B. Shirsath, Arshad Khan, Dhruv Kumar, Balvinder K. Sapra

## Abstract

**Purpose:** Low-dose radiation-induced adaptive response (LDRIAR) is well documented, but its role in early DNA damage signalling remains unclear. This study aimed to investigate whether adaptive response influences initial DNA double-strand break (DSB) recognition, as reflected by γH2AX foci formation, and to evaluate its time-dependent expression in human lymphocytes.

**Materials and Methods:** Peripheral blood lymphocytes from three healthy donors were exposed to a priming dose followed by a challenging dose at defined time intervals. DNA damage was assessed using γH2AX foci analysis, comparing acute and split-dose exposures in both PHA-stimulated (large) and non-stimulated (small) lymphocytes.

**Results:** A clear time-dependent adaptive response was observed. No significant reduction in γH2AX foci was detected at 1 h (p > 0.05). At 2 h, a significant decrease was observed (∼7–8% in large and ∼13% in small lymphocytes; p < 0.01), which increased at 4 h (∼12% and ∼22%, respectively; p < 0.001). The maximal response occurred at 15 h, with reductions of ∼40– 43% in large and ∼27% in small lymphocytes (p < 0.001). Small lymphocytes exhibited an earlier response, while large lymphocytes showed a greater magnitude at later time points. The temporal trend was consistent across donors, with minor variability at later intervals.

**Conclusions:** The findings demonstrate that LDRIAR is reflected at the level of DNA damage signalling and follows a defined temporal pattern with cell-type specificity. This suggests that adaptive response may influence early DSB-associated processes, contributing to a better understanding of radiation response mechanisms in radiobiology.

## 1. Introduction

Ionizing radiation (IR) interacts with biological systems in diverse and complex ways, producing outcomes that depend on multiple factors including radiation dose, dose rate, linear energy transfer (LET), exposure history, and intrinsic biological susceptibility. Conventionally, the biological effects of radiation have been assumed to follow a linear dose-response relationship with no safe threshold, forming the basis of the widely applied Linear No-Threshold (LNT) model for radiation protection (Tapio and Jacob 2007). However, growing experimental evidence suggests that low-dose ionizing radiation (LDIR), typically defined as <200 mGy, may elicit nonlinear biological responses distinct from those induced at high doses (Tapio and Jacob 2007; Tubiana et al. 2009; Chaurasia R and Sapra 2024). One of the consistently observed nonlinear responses is the low dose radiation-induced adaptive response (LDRIAR), wherein a small priming dose induces a protective cellular state that reduces toxic damage from a subsequent higher challenge dose.

The adaptive response was first demonstrated in human peripheral blood lymphocytes in 1984, when Olivieri et al. observed that cells pre-exposed to low concentrations of tritiated thymidine exhibited fewer chromosomal aberrations after subsequent X-irradiation compared to unprimed controls(Olivieri et al. 1984). This innovative finding challenged established radiobiological assumptions and stimulated extensive research into low-dose radiobiology. Follow-up studies in the late 1980s and early 1990s replicated the phenomenon in human lymphocytes, demonstrating dose-window specificity, time-dependent induction, and inter-donor variability (Sankaranarayanan et al. 1989; Feinendegen et al. 2004). Since then, RIAR has been documented in a broad range of biological systems including mammalian cells, plants, microorganisms, and in vivo animal models, indicating that it represents a conserved evolutionary stress response (Mortazavi et al. 2003; Guéguen et al. 2019).

Mechanistic investigations reveal that RIAR is not a single pathway but a coordinated biological program involving molecular, epigenetic, and metabolic reprogramming. Low-dose irradiation triggers mild oxidative stress and activates intracellular signaling pathways involving ATM, ATR, p53, MAPK, and NF-κB, subsequently modifying DNA repair fidelity, antioxidant capacity, apoptosis thresholds, and gene expression patterns (Szumiel 1998; Feinendegen et al. 2007; Guéguen et al. 2019). Enhanced activity of DNA repair pathways-particularly non-homologous end-joining (NHEJ) and base-excision repair (BER), has been reported as a major contributor to the observed reduction in chromosomal damage following challenge exposure (Sokolov et al. 2005). Similarly, transient upregulation of antioxidant enzymes such as glutathione peroxidase, catalase, and superoxide dismutase has been linked to reduced reactive oxygen species (ROS) burden during subsequent irradiation (Feinendegen et al. 2004).

Despite robust experimental documentation, the expression of adaptive response is highly variable. Its induction depends on several key parameters including priming dose (often 1-100 mGy), radiation type, dose rate, and the time interval between priming and challenge exposures, which typically ranges between 2 and 24 hours for optimal expression (Shadley and Wiencke 1989; Wolff 1998). Importantly, inter-individual variability has been repeatedly reported in human studies, suggesting contributions from genetic polymorphisms, age, immune status, oxidative stress baseline, and epigenetic factors (Tapio and Jacob 2007; Tang and Loke 2015; Lewis et al. 2017). Some studies have failed to observe RIAR entirely, emphasizing that it is conditional rather than universal (Fornalski et al. 2024; Kanani et al. 2025; Krasowska and Fornalski 2025). The lack of response under certain exposure conditions has contributed to continued discourse regarding its relevance to regulatory frameworks.

The implications of LDRIAR span multiple domains. From a radiation protection perspective, the existence of adaptive responses challenges the assumption that all low doses are biologically harmful and may support consideration of threshold or hormetic models in certain exposure contexts (Szumiel 1998; Feinendegen et al. 2004; Tubiana et al. 2009). In clinical radiobiology, adaptive response research has attracted interest for its potential therapeutic applications, particularly in reducing normal-tissue toxicity during radiotherapy or modifying radiosensitivity in personalized treatment planning. Preliminary studies suggest that low-dose pre-exposure may modulate inflammatory signaling and immune microenvironment, with possible implications for tumor control and normal-tissue sparing (Constanzo et al. 2021; Averbeck 2023).

With increasing exposure of humans to low-dose radiation through advanced medical imaging, space travel, nuclear industry expansion, and environmental sources, the biological significance of LDRIAR is increasingly relevant (Vaiserman et al. 2018; Pearson et al. 2021). Recent advances in high-throughput sequencing, single-cell profiling, and systems biology now enable comprehensive interrogation of adaptive response signatures, offering opportunities to refine mechanistic models and identify predictive biomarkers of radiation sensitivity and adaptive potential.

Taken together, LDRIAR remains one of the most intriguing and actively debated phenomena in low-dose radiobiology. Rigorous investigation using well-defined experimental systems, such as human peripheral blood lymphocytes, an established and physiologically relevant model, is essential to delineate the magnitude, inter-individual variability, and biological significance of this response.

In the present study, we systematically examine LDRIAR under controlled priming and challenge dose conditions, employing γH2AX foci formation as a quantitative surrogate of DNA-DSBs in PHA-stimulated human peripheral blood lymphocytes. Notably, while previous studies have predominantly linked LDRIAR to downstream cytogenetic endpoints reflecting misrepair, the current work addresses a critical and unresolved question: whether LDRIAR operates at the level of initial DNA-DSB induction itself or is restricted to modulation of repair fidelity. To the best of our knowledge, this represents the first study to directly interrogate the mechanistic basis of LDRIAR at the level of DSB induction using a γH2AX-based approach.

## 2. Materials and methods

### 2.1 Chemicals

Triton X-100, paraformaldehyde, and 1,1,3,3-tetraethoxypropane were obtained from Gibco Life Technologies (USA). Phytohemagglutinin (PHA), foetal calf serum (FCS), Hank’s Balanced Salt Solution (HBSS), and Roswell Park Memorial Institute medium (RPMI-1640) were also purchased from Gibco Life Technologies (USA). Penicillin and streptomycin were procured from HiMedia Laboratories (India). Mouse anti-phospho-histone H2AX (Ser139) monoclonal IgG (γH2AX) was obtained from Millipore (USA), and fluorescein isothiocyanate (FITC)-conjugated rabbit anti-mouse IgG secondary antibody was purchased from Invitrogen (USA). Nuclear counterstaining was performed using 4′,6-diamidino-2-phenylindole (DAPI) in an antifade mounting medium (ProLong™ Gold Antifade Mountant with DAPI; Invitrogen, USA).

### 2.2 Ethical approval and blood collection

Three healthy volunteers (Volunteer 1: male, 27 years; Volunteer 2: male, 41 years; Volunteer 3: female, 24 years) were recruited to investigate LDRIAR in peripheral blood lymphocytes. The study protocol was reviewed and approved by the Institutional Ethics Committee of the Bhabha Atomic Research Centre (Ref: BARCHMEC/59/2024), Mumbai, India. Written informed consent was obtained from all participants prior to sample collection. Peripheral blood (16 mL) was collected in heparinized vacutainers under aseptic conditions. All experimental procedures were conducted in strict accordance with the approved ethical guidelines and regulatory standards.

### 2.3 Isolation of peripheral blood lymphocytes and stimulation with PHA

Peripheral blood lymphocytes were isolated from whole blood by density gradient centrifugation using Histopaque-1077, with minor modifications to standard protocols (Dagur and McCoy 2015; Chaurasia RK et al. 2022). Briefly, blood samples were diluted 1:1 with HBSS and carefully layered onto the density gradient medium in sterile centrifuge tubes. Following centrifugation at 2,500 rpm for 25 min at room temperature, distinct phase separation was achieved. The mononuclear cell layer (buffy coat) was gently aspirated, washed twice with RPMI and resuspended at a final density of 1 × 10□ cells/mL. Lymphocyte suspension was divided into 12 equal aliquots (1 mL each) to facilitate parallel experimental treatments. For each condition, cultures were established in a total volume of 5 mL comprising 3.5 mL RPMI, 1 mL cell suspension, and 0.5 mL serum. Lymphocyte activation was induced by the addition of 0.1 mL PHA (Chaurasia Rajesh Kumar et al. 2025).

### 2.4 Irradiation

Out of twelve, four aliquots were designated for background and priming dose assessment. One sample was maintained as an unirradiated control to determine baseline γH2AX foci frequency. The remaining three samples were exposed to a priming dose of 0.03 Gy and incubated for 30 min, 1 h, and 2 h, respectively, to characterize the induction and resolution kinetics of DNA damage following low-dose exposure. The remaining eight aliquots were used to evaluate the LDRIAR. Of these, four samples served as corresponding acute exposure controls, while the other four were subjected to the split-dose protocol. For adaptive response induction, a priming dose of 0.03 Gy was delivered at 30 h post-PHA stimulation, followed by a challenging dose of 1.03 Gy administered at 1, 2, 4, and 15 h after priming. These time intervals correspond to 31, 32, 34, and 45 h post-PHA stimulation, respectively. Matched control samples were exposed to a single acute dose of 1.03 Gy at the corresponding time points (31, 32, 34, and 45 h post-PHA stimulation) to account for temporal variations in cellular response independent of priming. Similar experiments were conducted independently for two additional volunteers.

All irradiations were performed using a Blood Irradiator-2000 (BRIT, DAE, India). Distinct dose rates were employed for priming and challenging exposures to ensure controlled dose delivery: the priming dose (0.03 Gy) was delivered at 0.09 Gy/min, whereas the challenging dose (1.03 Gy) was delivered at 0.36 Gy/min. Irradiation conditions were standardized in accordance with established radiobiological protocols to ensure accuracy and reproducibility. Samples were irradiated in a water-equivalent phantom with a 3 mm buildup layer to maintain electronic equilibrium, and both phantom and samples were pre-equilibrated to 37°C to approximate physiological conditions. All irradiations for each donor were completed within a single day under identical conditions to minimize experimental variability. Dosimetry was performed using a Fricke dosimeter calibrated to national standards traceable to international references, ensuring accurate dose delivery (Chaurasia R. K. et al. 2025).

### 2.5 γH2AX Immunofluorescence staining and foci analysis

To arrest DNA repair, irradiated lymphocytes were immediately placed on ice and subsequently incubated at 37 °C (5% CO□) for 30 min to permit H2AX phosphorylation. Immunofluorescence staining was performed using a laboratory-optimized protocol in accordance with the antibody supplier’s guidelines (Chaurasia RK et al. 2021). Cells were then fixed in chilled 4% paraformaldehyde (in PBS) for 30 min at 4 °C, followed by three PBS washes (5 min each). Permeabilization was performed using 0.5% Triton X-100 in PBS for 3 min, followed by two PBS washes (5 min each). Cells (100 µL) were seeded onto poly-L-lysine-coated 22 × 22 mm coverslips placed in 35 mm Petri dishes and allowed to adhere. Blocking was carried out with 5% FCS in PBS for 1 h without subsequent washing. Cells were incubated with mouse monoclonal anti-γH2AX primary antibody (1:400) for 1 h at 37 °C in a humidified chamber, followed by washing (two PBS washes and one PBST wash, PBS + 0.1% Tween-20). Secondary staining was performed using Alexa Fluor 488-conjugated rabbit anti-mouse antibody (1:600) for 45 min at 37 °C, followed by identical washing steps. Nuclei were counterstained and mounted using DAPI-containing antifade medium, incubated for 10 min at room temperature, and sealed. γH2AX foci were visualized using a Leica DM8 confocal microscope with appropriate filter settings. A minimum of 100 cells per sample were analyzed to determine foci frequency per nucleus for each dose and time point.

### 2.6 Statistical analysis

Initial comparisons between groups were performed using a two-tailed Z-test, with variance estimated from the corresponding SDs, and percentage change in foci yield was calculated to assess the magnitude of response. For cumulative analysis across three independent volunteers, SDs were pooled using a weighted variance approach assuming equal sample size per group. To account for inter-individual variability and repeated measurements, a linear mixed-effects model was applied with dose and time as fixed effects and volunteer as a random effect, including their interaction (Delafont et al. 2018; Liu et al. 2024). Where appropriate, post hoc comparisons were performed at individual time points. Statistical significance was defined at p < 0.05. Z-values were not included in Table 4 (cumulative data of all three volunteers), as final statistical inferences were based on the mixed-effects model, which accounts for inter-individual variability and repeated measures (Delafont et al. 2018; He and Lee 2022).

## 3 Results

### 3.1 Time-dependent induction of adaptive response following split-dose irradiation

The induction of LDRIAR was evaluated by comparing γH2AX foci yields between split-dose (priming + challenging; 0.03 + 1.03 Gy) and single acute dose (1.03 Gy) exposures at multiple time intervals following priming. It is important to note that PHA stimulation resulted in activation of only ∼40-65% of the total lymphocyte population (Lkhagvasuren 2017; Abbas et al. 2021). The large lymphocyte (diameter ≥ 10 µm) population predominantly represents PHA-stimulated (activated) cells, whereas the small lymphocyte (diameter ≤ 8 µm) population corresponds to non-stimulated (resting) cells (Figure 1) (Tigner et al. 2020).

**Figure 1.**
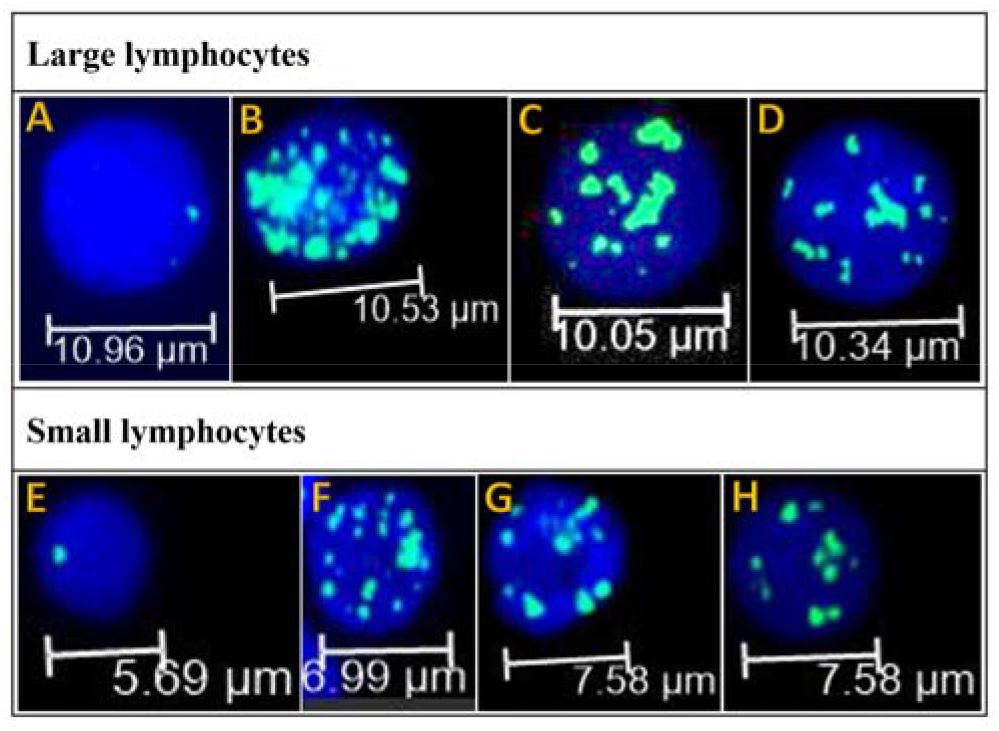
Representative peripheral blood lymphocytes showing γH2AX foci (green) with nuclei counterstained by DAPI (blue). Both PHA-stimulated large lymphocytes and non-stimulated small lymphocytes are shown. (A–D) PHA-stimulated large cells: (A) background (basal) foci; (B–D) γH2AX foci at 2, 4, and 15 h after the priming dose, respectively. (E–H) Non-stimulated small lymphocytes: (E) background foci; (F–H) γH2AX foci at 2, 4, and 15 h after the priming dose, respectively. A progressive reduction in foci number with increasing post-priming time is observed in both cell types, indicating an adaptive response at the level of DNA-DSB formation.

Across all three volunteers, a clear time-dependent modulation of DNA damage response was observed, characterized by a progressive reduction in γH2AX foci in the split-dose condition relative to the acute exposure. At 1 h post-priming, no significant differences in foci yield were detected between the two exposure conditions in either large or small lymphocyte populations (p > 0.05), indicating the absence of a measurable adaptive response at this early time point. However, from 2 h onwards, a statistically significant reduction in foci yield emerged (Table 1, 2 and 3), demonstrating the onset of adaptive response, which further intensified at later time points (Figure 2, 3 and 4).

**Table 1.**
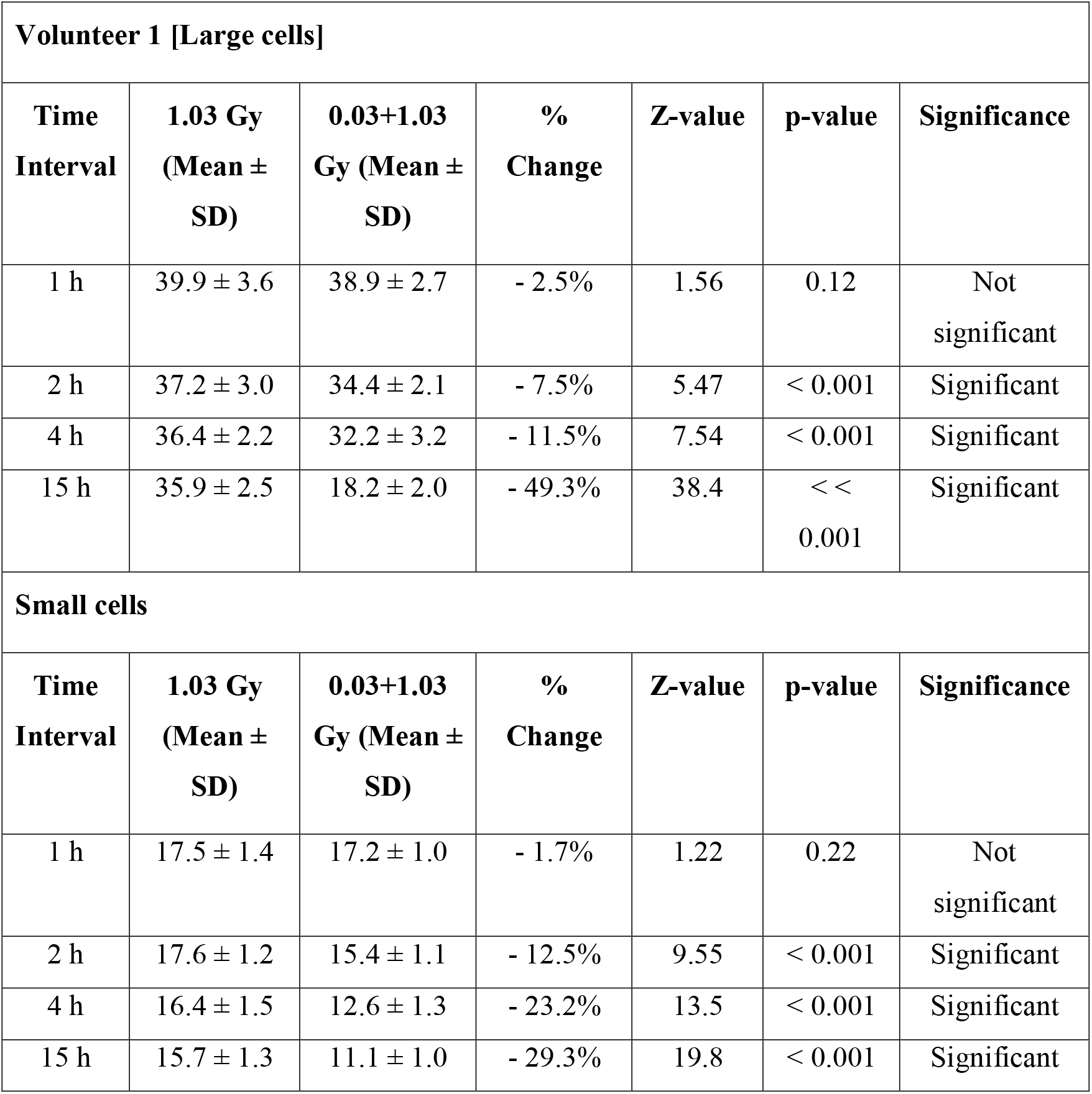
Time-dependent comparison of DNA damage response ((γH2AX foci)) in large and small lymphocyte populations from Volunteer 1 following acute (1.03 Gy) and split-dose (0.03 + 1.03 Gy) irradiation. Values are expressed as mean ± SD at indicated time points. Percentage change denotes variation in the split-dose relative to acute exposure. Statistical significance between groups was assessed using Z-test; p < 0.05 was considered significant.

| Volunteer 1 [Large cells] |  |  |  |  |  |  |
| --- | --- | --- | --- | --- | --- | --- |
| Time Interval | 1.03 Gy (Mean ± SD) | 0.03+1.03 Gy (Mean ± SD) | % Change | Z-value | p-value | Significance |
| 1 h | 39.9 ± 3.6 | 38.9 ± 2.7 | - 2.5% | 1.56 | 0.12 | Not significant |
| 2 h | 37.2 ± 3.0 | 34.4 ± 2.1 | - 7.5% | 5.47 | < 0.001 | Significant |
| 4 h | 36.4 ± 2.2 | 32.2 ± 3.2 | - 11.5% | 7.54 | < 0.001 | Significant |
| 15 h | 35.9 ± 2.5 | 18.2 ± 2.0 | - 49.3% | 38.4 | < < 0.001 | Significant |
| Small cells |  |  |  |  |  |  |
| Time Interval | 1.03 Gy (Mean ± SD) | 0.03+1.03 Gy (Mean ± SD) | % Change | Z-value | p-value | Significance |
| 1 h | 17.5 ± 1.4 | 17.2 ± 1.0 | - 1.7% | 1.22 | 0.22 | Not significant |
| 2 h | 17.6 ± 1.2 | 15.4 ± 1.1 | - 12.5% | 9.55 | < 0.001 | Significant |
| 4 h | 16.4 ± 1.5 | 12.6 ± 1.3 | - 23.2% | 13.5 | < 0.001 | Significant |
| 15 h | 15.7 ± 1.3 | 11.1 ± 1.0 | - 29.3% | 19.8 | < 0.001 | Significant |

**Table 2.**
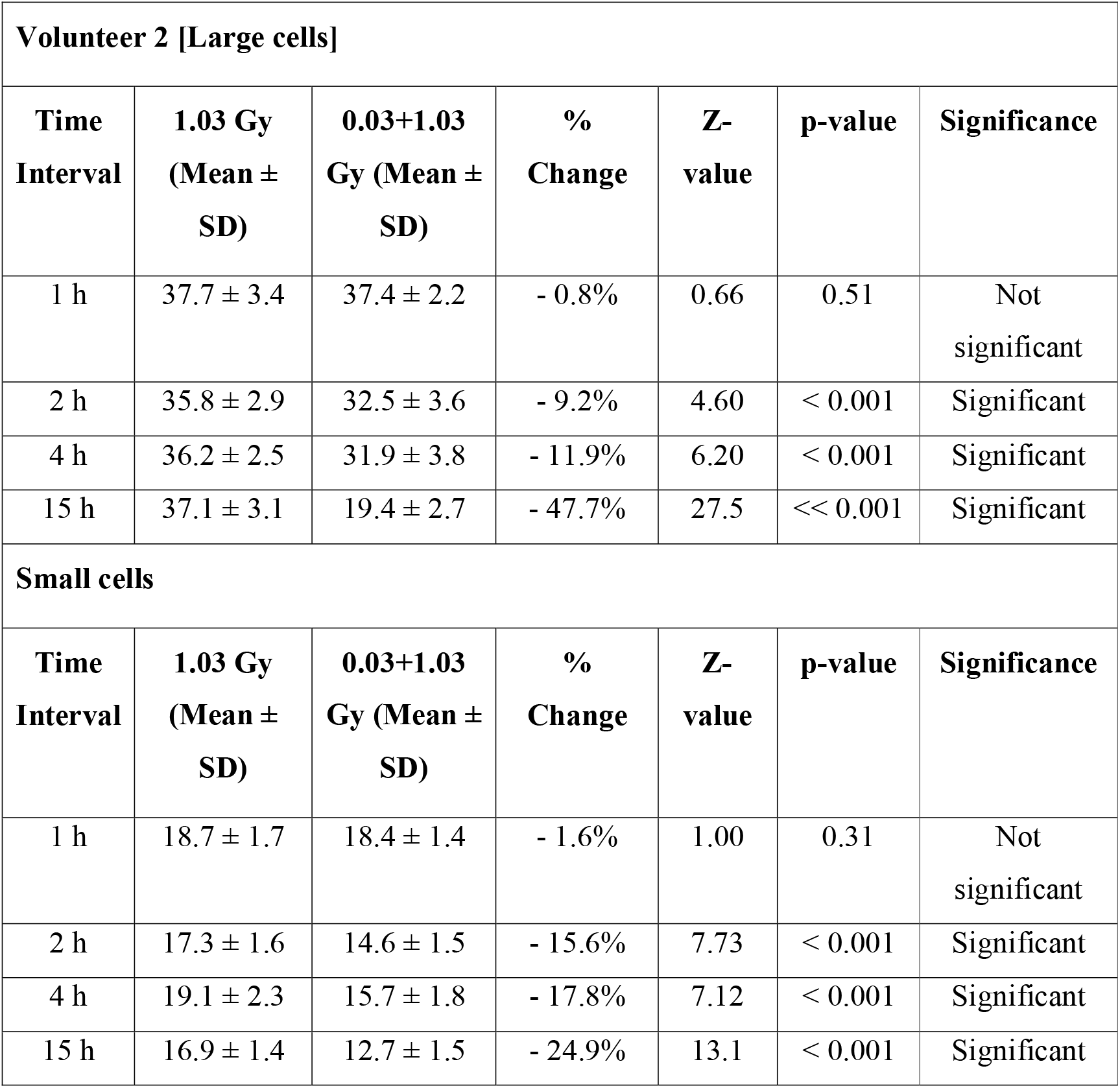
Time-dependent comparison of DNA damage response (γH2AX foci) in large and small lymphocyte populations from Volunteer 2 following acute (1.03 Gy) and split-dose (0.03 + 1.03 Gy) irradiation. Values are expressed as mean ± SD. Percentage change represents variation in the split-dose relative to acute exposure. Statistical analysis was performed using Z-test; p < 0.05 was considered significant.

**Table 3.** Time-dependent comparison of DNA damage response (γH2AX foci) in large and small lymphocyte populations from Volunteer 3 following acute (1.1 Gy) and split-dose (0.1 + 1 Gy) irradiation. Values are presented as mean ± SD. Percentage change indicates variation in the split-dose relative to acute exposure. Statistical significance was evaluated using Z-test; p < 0.05 was considered significant.

| **Volunteer 3 [Large cells]** |  |  |  |  |  |  |
| --- | --- | --- | --- | --- | --- | --- |
| **Time Interval** | **1.03 Gy (Mean ± SD)** | **0.03+1.03 Gy (Mean ± SD)** | **% Change** | **Z-value** | **p-value** | **Significance** |
| 1 h | 38.3 ± 3.0 | 38.7 ± 2.5 | + 1.0% | 0.74 | 0.46 | Not significant |
| 2 h | 40.1 ± 3.6 | 37.4 ± 2.8 | - 6.7% | 3.83 | < 0.001 | Significant |
| 4 h | 37.8 ± 2.9 | 32.7 ± 3.3 | - 13.5% | 7.49 | < 0.001 | Significant |
| 15 h | 38.3 ± 3.1 | 26.4 ± 2.6 | -31.1% | 20.8 | < 0.001 | Significant |
| **Small cells** |  |  |  |  |  |  |
| **Time Interval** | **1.03 Gy (Mean ± SD)** | **0.03+1.03 Gy (Mean ± SD)** | **% Change** | **Z-value** | **p-value** | **Significance** |
| 1 h | 18.2 ± 1.5 | 17.9 ± 1.3 | - 1.6% | 1.07 | 0.28 | Not significant |
| 2 h | 20.1 ± 1.8 | 17.9 ± 1.5 | - 10.9% | 5.77 | < 0.001 | Significant |
| 4 h | 16.9 ± 1.4 | 12.6 ± 1.5 | - 25.4% | 14.5 | < 0.001 | Significant |
| 15 h | 18.2 ± 1.6 | 13.2 ± 1.4 | - 27.5% | 15.1 | < 0.001 | Significant |

**Figure 2.**
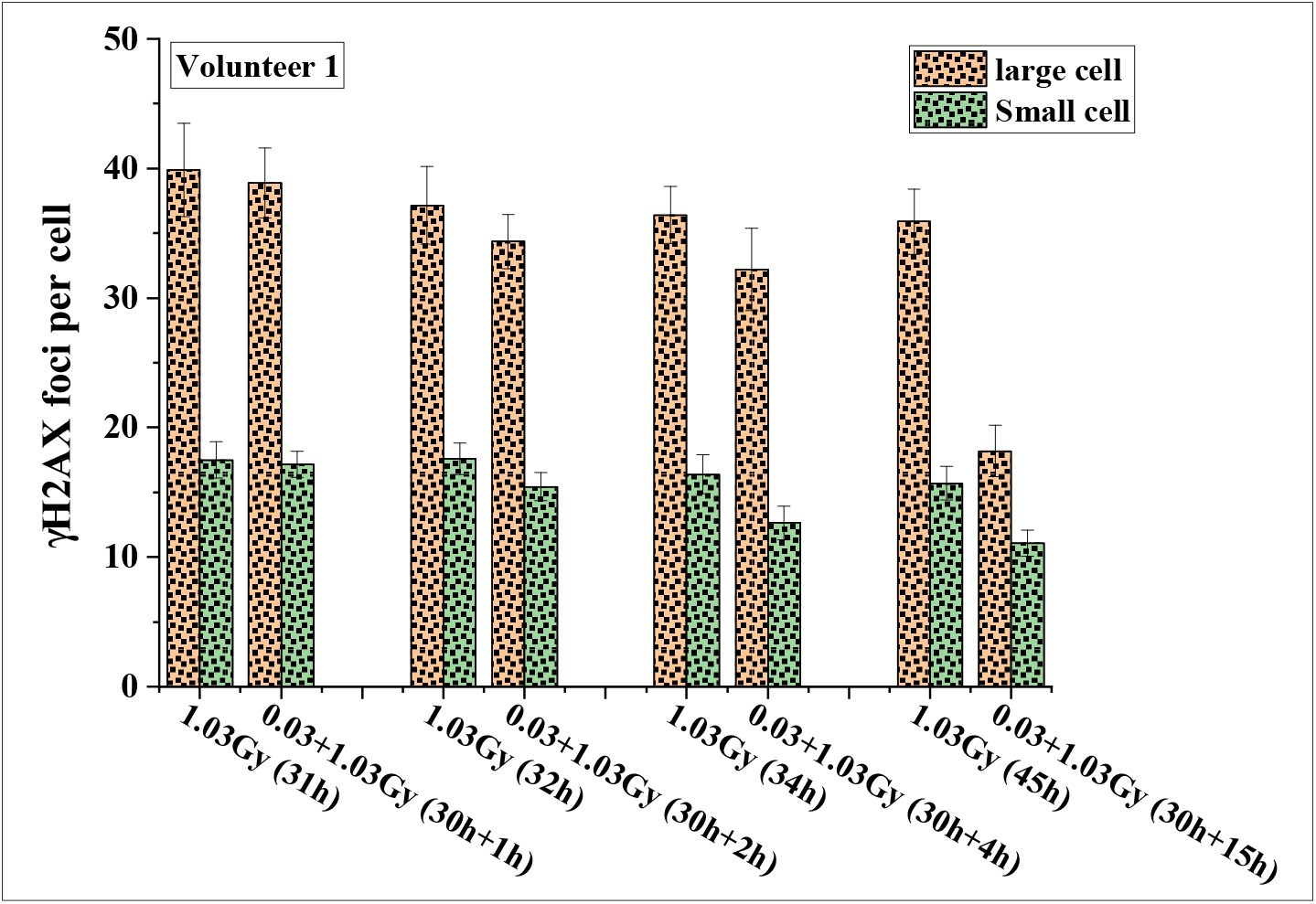
Time-dependent variation in γH2AX foci per cell in large and small lymphocyte populations from Volunteer 1 following acute (1.03 Gy) and split-dose (0.03 + 1.03 Gy) irradiation. Data represent mean ± SD at indicated post-irradiation time points. Split-dose exposure shows a progressive reduction in foci compared to acute dose, more pronounced at later time points.

**Figure 3.**
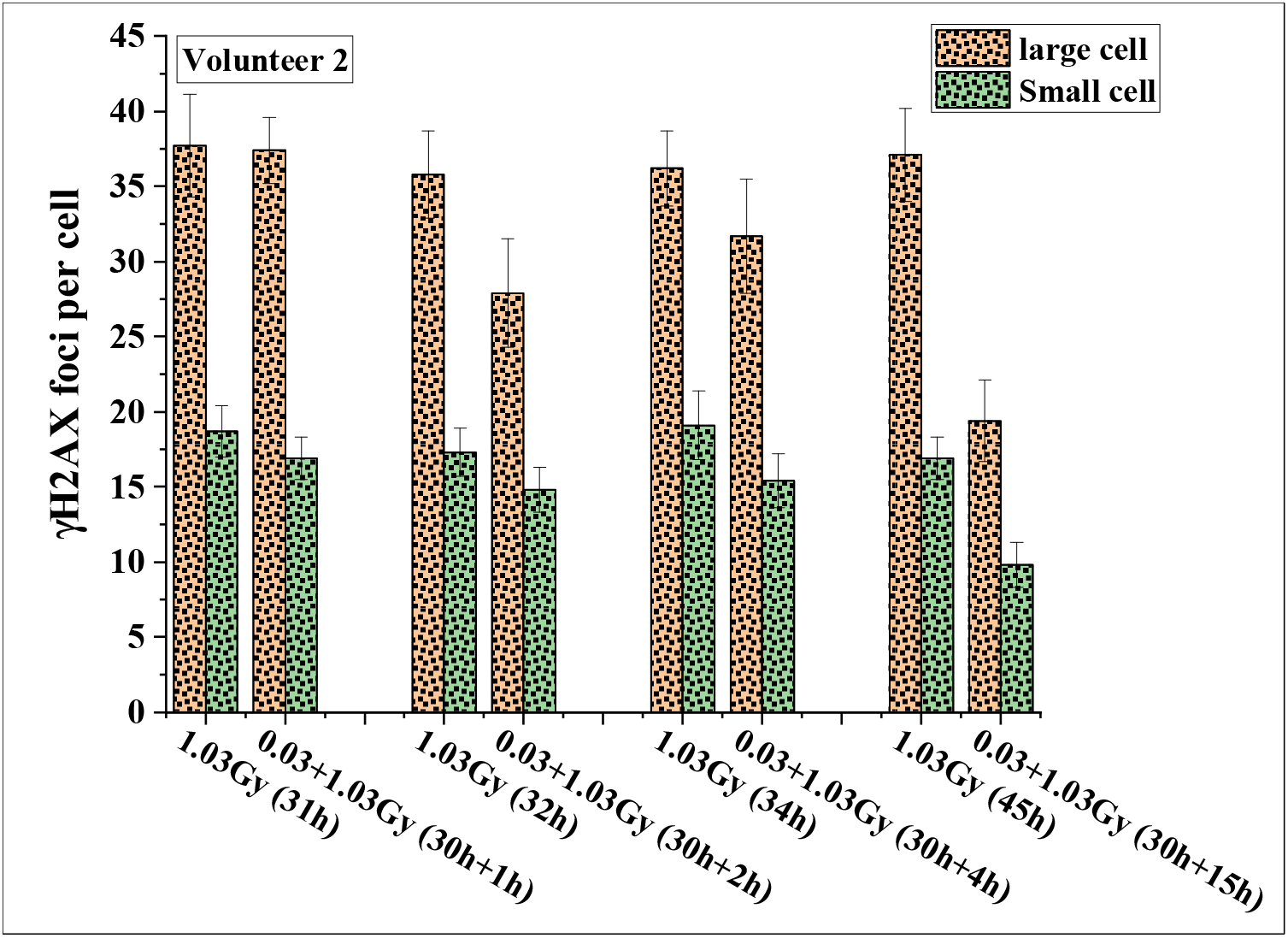
Time-dependent variation in γH2AX foci per cell in large and small lymphocyte populations from Volunteer 2 following acute (1.03 Gy) and split-dose (0.03 + 1.03 Gy) irradiation. Values are presented as mean ± SD at indicated time points. A progressive reduction in foci is observed in the split-dose condition compared to acute exposure, particularly at later time points.

**Figure 4.**
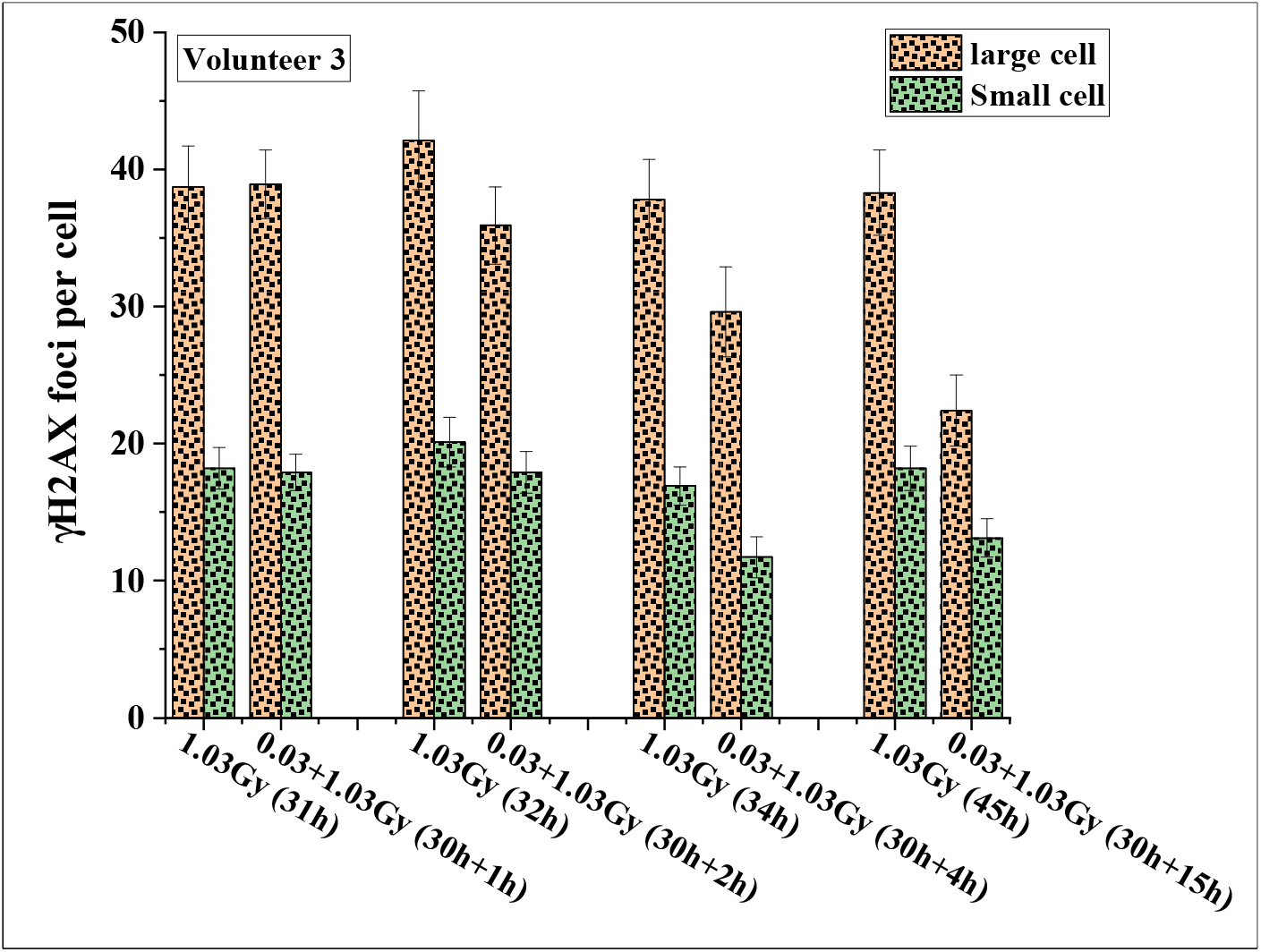
Time-dependent variation in γH2AX foci per cell in large and small lymphocyte populations from Volunteer 3 following acute (1.03 Gy) and split-dose (0.03 + 1.03 Gy) irradiation. Data are expressed as mean ± SD at indicated time points. A reduction in foci is evident in the split-dose condition relative to acute exposure, with a more pronounced effect at later time points.

Baseline γH2AX foci in human lymphocytes (pooled across three volunteers) ranged from ∼0.1-0.6 per cell. Following exposure to 0.03 Gy (priming dose), a transient increase to ∼0.3-0.9 foci per cell was observed at 30 min, which declined to ∼0.2-0.75 at 1 h and ∼ 0.15-0.7 at 2 h. This temporal pattern is consistent with rapid double-strand break repair kinetics (Horn et al. 2011; Chaurasia RK et al. 2021). No statistically significant differences from controls were detected at any time point (0.5, 1, and 2 h; p > 0.05). Foci levels returned to baseline by 2 h, indicating complete resolution with no residual effect at the time of the subsequent challenging dose. All values presented in Tables 1-3 are background-corrected.

### 3.2 Differential responses in stimulated and non-stimulated lymphocytes

A distinct population-dependent variation in adaptive response was observed. In large lymphocytes, the reduction in γH2AX foci following split-dose exposure was modest at 2 h (∼6-9%) and increased further at 4 h (∼ 11-13%), culminating in a marked decrease at 15 h (∼31-49%) across all three volunteers (p < 0.001).

In contrast, small lymphocytes exhibited a comparatively attenuated response. Although a significant reduction in foci yield was evident from 2 h (∼10-16%) and 4 h (∼17-25%) onwards, the magnitude of reduction at 15 h (∼24-29%) remained consistently lower than that observed in large lymphocytes. These findings indicate a more pronounced adaptive response in the large lymphocyte population, particularly at extended time intervals following priming.

### 3.3 Inter-individual variability and consistency of adaptive response

While some degree of inter-individual variability was evident in the magnitude of response, the overall temporal pattern of LDRIAR induction was consistent across all three volunteers. Each donor demonstrated negligible response at 1 h, followed by a significant and progressive reduction in foci yield at 2 h and 4 h, and a maximal response at 15 h.

This consistency across independent biological samples underscores the reproducibility of the observed adaptive response despite inherent biological variability.

### 3.4 Cumulative analysis across volunteers

To further validate these observations, cumulative data pooled from all three volunteers were analyzed (Table 4, Figure 5). In addition, inter-individual variability and differences between lymphocyte subpopulations were evaluated. The pooled analysis reinforced the time-dependent nature of the adaptive response. At 1 h post-priming, no statistically significant reduction in γH2AX foci was observed in either large or small lymphocytes (p > 0.05), and importantly, no substantial inter-individual variation was apparent among the three volunteers at this early time point, indicating a consistent lack of adaptive response initiation. At 2 h, a statistically significant reduction in foci was observed in both large (∼7.8%) and small (∼13.0%) lymphocytes (p < 0.01). While all three volunteers exhibited a similar trend, minor differences in magnitude were noted; however, these variations were not markedly different across volunteers, indicating that the adaptive response at this stage is consistent across individuals. Notably, the reduction was greater in small lymphocytes compared to large lymphocytes, suggesting an earlier or more sensitive response in the small cell population. At 4 h, the reduction became more pronounced, with decreases of ∼12.3% in large cells and ∼22.1% in small cells (p < 0.001). Inter-individual differences remained modest, confirming reproducibility across donors. However, the difference between lymphocyte subpopulations was evident, with small lymphocytes consistently showing a greater reduction in foci than large lymphocytes. At 15 h, the maximal adaptive response was observed, with a substantial reduction of ∼42.6% in large lymphocytes and ∼27.2% in small lymphocytes (p < 0.001). At this time point, although some variability in magnitude persisted among volunteers, significant inter-individual variation was observed, with Volunteer 3 exhibiting a significantly weaker adaptive response compared to Volunteers 1 and 2 (p < 0.001), indicating that the adaptive response at later time points may vary between individuals. In contrast to earlier time points, the magnitude of reduction was greater in large lymphocytes compared to small lymphocytes, demonstrating a shift in dominance of adaptive response toward the large (PHA-stimulated) cell population at later times.}

**Table 4.**
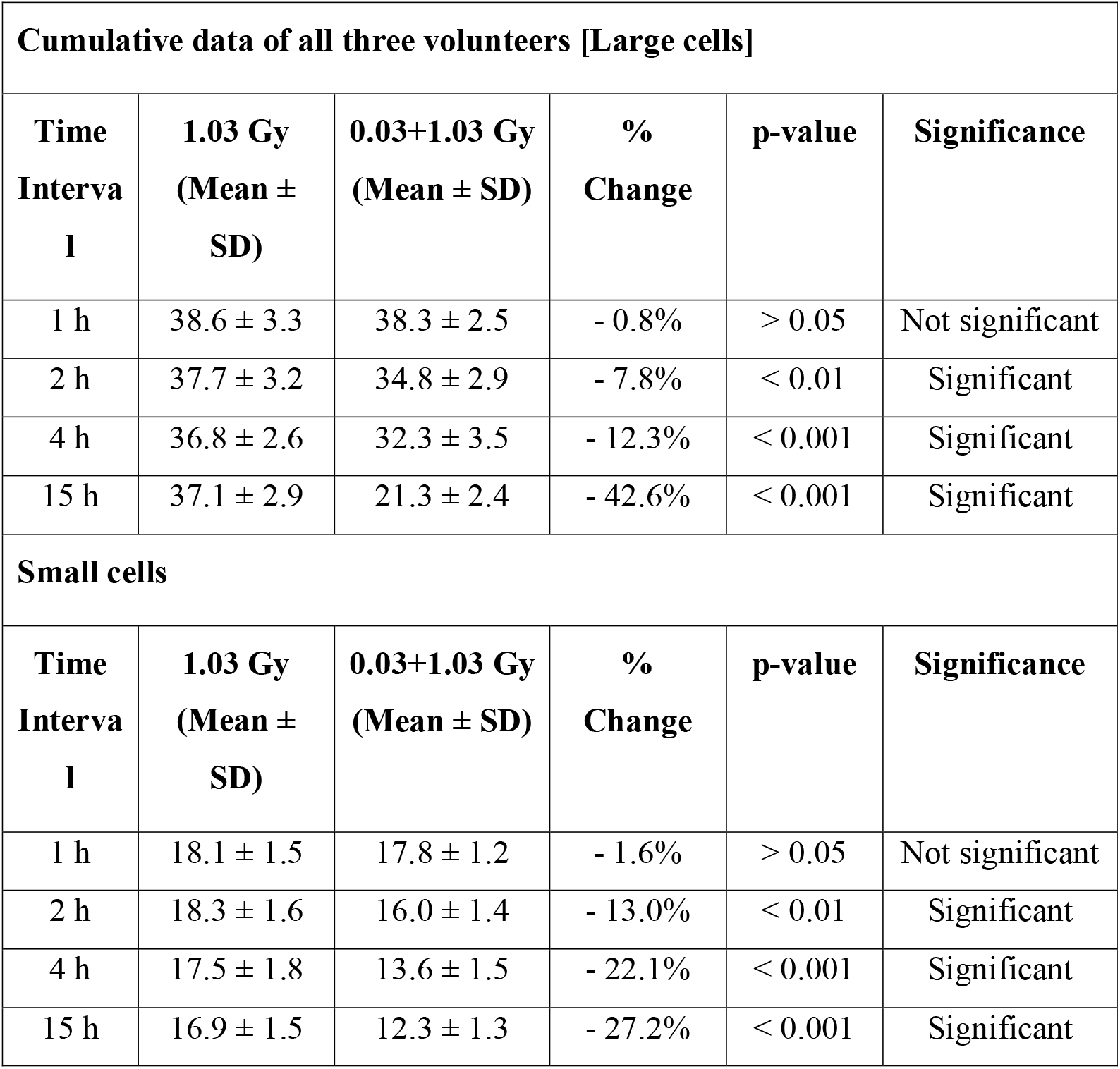
Cumulative analysis of DNA damage response (γH2AX foci) in large and small lymphocyte populations pooled from three volunteers following acute (1.03 Gy) and split-dose (0.03 + 1.03 Gy) irradiation. Values are expressed as mean ± SD at indicated time points. Percentage change represents variation in the split-dose relative to acute exposure. Statistical significance between groups was determined using appropriate comparative analysis; p < 0.05 was considered significant.

**Figure 5.**
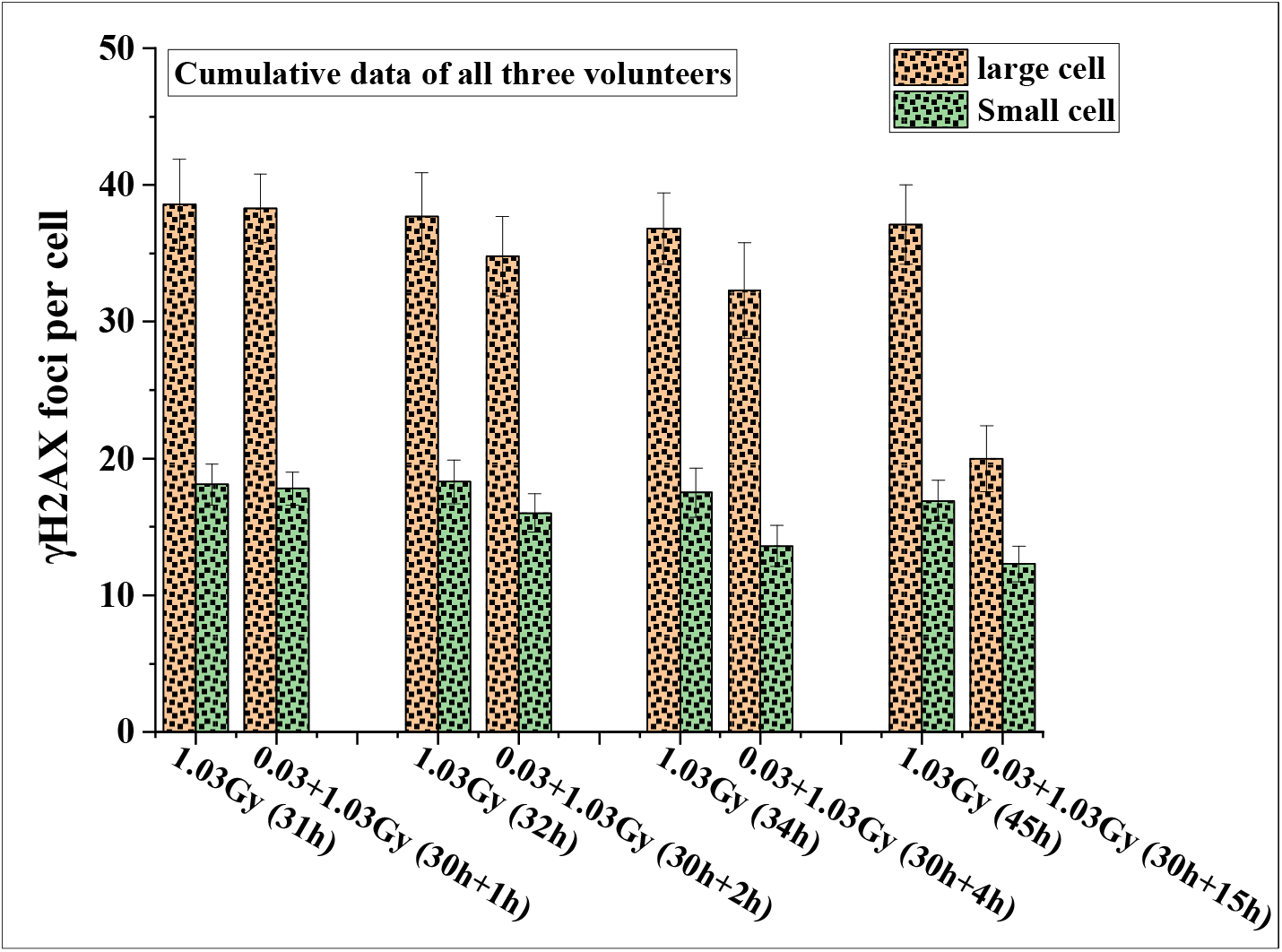
Cumulative analysis of γH2AX foci per cell in large and small lymphocyte populations pooled from all three volunteers following acute (1.03 Gy) and split-dose (0.03 + 1.03 Gy) irradiation. Values are presented as mean ± SD at indicated time points. The split-dose condition shows a consistent reduction in foci compared to acute exposure, with a more pronounced effect at later time points.

### 3.5 Overall trend and magnitude of adaptive response

Collectively, the data demonstrate a clear and progressive enhancement of LDRIAR with increasing time interval between priming and challenging doses. The absence of response at early time points, followed by a gradual and significant reduction in DNA damage at later intervals, indicates that the development of adaptive response is a time-dependent process. Importantly, the magnitude of the adaptive response was generally greater in PHA-stimulated (large) lymphocytes compared to non-stimulated (small) lymphocytes, particularly at later time points, suggesting differences in cellular activation status, cell cycle progression, or DNA repair capacity between these populations.

## Discussion

The present study provides direct experimental evidence that LDRIAR manifests at the level of DNA DSB signalling, as quantified by γH2AX foci, rather than being restricted solely to downstream cytogenetic endpoints. The findings demonstrate a clear time-dependent reduction in γH2AX foci following split-dose irradiation, with statistically significant effects emerging beyond 2 h post-priming and reaching maximal magnitude at 15 h. Importantly, this response was observed across all three donors and exhibited distinct differences between PHA-stimulated (large) and non-stimulated (small) lymphocyte populations.

A key observation of this study is the absence of any measurable adaptive response at 1 h post-priming, followed by a progressive and statistically significant reduction in γH2AX foci at 2 h, 4 h, and 15 h. This temporal pattern strongly supports the notion that LDRIAR is not an immediate phenomenon but requires a finite induction period. Similar time-dependent induction windows have been reported in earlier studies employing cytogenetic endpoints (representing post-repair events) where optimal adaptive responses were observed within 2-24 h following priming exposure in unstimulated lymphocytes (Wolff 1998; Feinendegen 2016). Mechanistically, this delay likely reflects the time required for activation of cellular signalling cascades triggered by low-dose radiation. Pathways involving ATM, p53, NF-κB, and MAPK are known to be activated following low-dose exposure and regulate downstream processes such as DNA repair, antioxidant defenses, and cell cycle control (Tapio and Jacob 2007; Azzam et al. 2012). The lack of response at 1 h suggests that these pathways have not yet reached sufficient activity to influence the cellular response to the subsequent challenge dose. The pronounced reduction in γH2AX foci at 15 h (∼43% in large lymphocytes, pooled data) further indicates that the adaptive response strengthens over time, likely reflecting transcription-dependent upregulation of DNA repair proteins and redox-modulating enzymes, as reported in low-dose radiation studies (Feinendegen et al. 2009).

A particularly important aspect of this study is the demonstration that adaptive response may influence γH2AX foci formation itself, a surrogate marker of initial DSB induction. Traditionally, LDRIAR has been interpreted as a consequence of improved DNA repair fidelity rather than reduced initial damage (Olivieri et al. 1984; Shadley and Wiencke 1989). However, the observed reduction in foci at early post-challenge time points suggests that priming may also modulate either the extent of initial DSB formation, possibly through chromatin remodelling or altered radiation track interactions, or the kinetics of very early DSB recognition and signalling, leading to reduced γH2AX accumulation. While γH2AX is widely used as a marker of DSBs, it reflects both damage induction and early signalling events (Rogakou et al. 1998). Therefore, the reduced foci observed here may indicate faster repair initiation or altered chromatin accessibility rather than a true decrease in physical DNA breakage. This interpretation remains speculative and warrants further investigation using complementary endpoints such as pulsed-field gel electrophoresis.

The study indicates a population-dependent variation in adaptive response. Small lymphocytes exhibited an earlier and relatively greater reduction at intermediate time points (2-4 h), whereas large (PHA-stimulated) lymphocytes demonstrated a more pronounced response at 15 h. This differential behavior can be explained by differences in cellular activation status and cell cycle distribution. PHA-stimulated lymphocytes are actively cycling and exhibit enhanced DNA repair capacity, particularly through homologous recombination and high-fidelity repair pathways (Jeggo and Löbrich 2006). In contrast, resting lymphocytes rely predominantly on non-homologous end joining (NHEJ), which is faster but more error-prone. The earlier response in small lymphocytes may reflect rapid activation of stress-response pathways, whereas the delayed but stronger response in large lymphocytes likely arises from transcription-dependent processes requiring active metabolism and cell cycle progression. This interpretation is consistent with previous reports showing that proliferating cells exhibit more robust adaptive responses due to greater plasticity in gene expression and repair pathway utilization (Sasaki 2003).

Although minor variations in magnitude were observed across donors, the overall temporal pattern of adaptive response remained consistent. However, at later time points (15 h), inter-individual variability became more evident, with one donor (Volunteer 3) exhibiting a comparatively weaker adaptive response than the others, which was statistically significant (p < 0.001, pairwise comparison with Volunteers 1 and 2). This observation suggests that while the general adaptive response trend is reproducible, its magnitude may vary between individuals. Such variability is consistent with previous reports highlighting donor-dependent differences in adaptive response (Bosi and Olivieri 1989; Scott 2014). Nonetheless, the limited sample size restricts definitive conclusions regarding population-level variability.

The magnitude of foci reduction observed, particularly the ∼31-49% decrease in large lymphocytes at 15 h across individual donors (and ∼42.6% in pooled analysis), is biologically substantial and suggests a meaningful enhancement of cellular resistance to radiation-induced DNA damage. These findings support the concept that low-dose exposure can induce a protective state, contributing to ongoing discussions regarding the biological effects of low-dose radiation and the assumptions underlying the LNT model (Feinendegen et al. 2004; Feinendegen et al. 2007; Tubiana et al. 2009).

Future studies may further strengthen and extend these findings by incorporating larger and more diverse donor cohorts to better characterize inter-individual variability, particularly at later time points. The inclusion of additional and complementary endpoints beyond γH2AX foci would provide a more comprehensive understanding of DNA damage induction and repair dynamics. Moreover, integrating direct mechanistic approaches, such as analyses of DNA repair pathway activity, chromatin organization, and signalling cascades, may help to elucidate the underlying biological processes governing the adaptive response. Such efforts would contribute to a more detailed and mechanistic understanding of LDRIAR and its potential implications in radiobiology.

## Conclusion

The present study demonstrates that low-dose radiation-induced adaptive response (LDRIAR) is expressed at the level of DNA double-strand break signalling, as reflected by γH2AX foci modulation. The findings establish a clear time-dependent induction pattern, with no measurable response at early time points and progressively enhanced protection at later intervals following priming. Importantly, the response exhibited cell-type specificity, with non-stimulated lymphocytes showing an earlier reduction in damage, whereas PHA-stimulated lymphocytes displayed a more pronounced effect at extended time intervals. This work provides novel evidence that adaptive response may influence early DNA damage signalling dynamics, extending beyond its traditionally understood role in post-damage repair processes. These observations have potential implications for understanding radiation response mechanisms and may inform strategies in radiobiology and radiation protection.

However, the study is limited by a small sample size and observed inter-individual variability at later time points, which may influence the generalizability of the findings. Future investigations incorporating larger cohorts and complementary mechanistic endpoints are required to further delineate the underlying processes and biological significance of LDRIAR.

## Data availability

All data generated in this study are included in the article. Additional data supporting the findings are available from the corresponding author (Email ID), upon reasonable request.

## Acknowledgement

The authors would like to express their sincere gratitude to Mr. Shrikant Jagtap, and other technical staff from our lab for their valuable assistance and technical support.

## Author information

### Authors and Affiliations

School of Health Sciences and Technology, University of Petroleum and Energy Studies, Dehradun, India

Sheeri Fatima and Dhruv kumar

Radiological Physics and Advisory Division, Bhabha Atomic Research Centre (BARC), Mumbai, India.

Rajesh Kumar Chaurasia, Aarti Notnani, K.B. Shirsath, Arshad Khan and B.K. Sapra

Homi Bhabha National Institute (HBNI), Mumbai, India.

B.K. Sapra

## Contributions

Rajesh Kumar Chaurasia, Sheeri Fatima and Aarti Notnani Aarti: Conceptualization, Methodology and Writing; Sheeri Fatima and Aarti Notnani and K.B. Shirsath: Methodology and Data Curation; B.K. Sapra, Arshad Khan and Dhruv kumar: Review, Editing, Supervision and Resources.

## Corresponding author

Correspondence to Dr. Rajesh Kumar Chaurasia

## Ethics declarations

### Ethics approval and consent to participate

The study protocol was reviewed and approved by the Institutional Ethics Committee of the Bhabha Atomic Research Centre (Ref: BARCHMEC/59/2024), Mumbai, India. All procedures involving human participants were conducted in accordance with the ethical standards of the committee. Written informed consent was obtained from all participants prior to their inclusion in the study.

### Consent for publication

Not applicable.

### Declaration of generative AI use

The authors confirm that they have read and complied with the Taylor & Francis AI Policy. During the preparation of this manuscript, the generative AI tool ChatGPT (OpenAI, GPT-5.3 version) was used solely for language correction and improvement of clarity. The tool assisted in refining grammar, sentence structure, and readability, while all scientific content, interpretation, and conclusions remain the responsibility of the authors.

## References

Abbas AK, Lichtman AH, Pillai S. 2021. Cellular and Molecular Immunology, 10e, South Asia Edition-E-Book. Elsevier Health Sciences.

Averbeck D. 2023. Low-dose non-targeted effects and mitochondrial control. International journal of molecular sciences. 24(14):11460.

Azzam EI, Jay-Gerin JP, Pain D. 2012. Ionizing radiation-induced metabolic oxidative stress and prolonged cell injury. Cancer letters. 327(1–2):48–60. eng.

Bosi A, Olivieri G. 1989. Variability of the adaptive response to ionizing radiations in humans. Mutation research. 211(1):13–17. eng.

Chaurasia R, Sapra B. 2024. Effects of Low Dose Ionizing Radiation on Human Health: Evidence for Revisiting Radiation Protection Policies. Handbook on Radiation Environment, Volume 1: Sources, Applications and Policies. Springer; p. 417–442.

Chaurasia RK, Bhat NN, Gaur N, Shirsath KB, Desai UN, Sapra BK. 2021. Establishment and multiparametric-cytogenetic validation of (60)Co-gamma-ray induced, phospho-gamma-H2AX calibration curve for rapid biodosimetry and triage management during radiological emergencies. Mutation research Genetic toxicology and environmental mutagenesis. 866:503354. eng.

Chaurasia RK, Pathak RS, Goel A, Shirsath KB, Bhat NN, Khan A, Sapra BK. 2025. First evidence of coexistence of Pseudo Pelger Huet anomaly and balanced translocation in a two decades retrospectively exposed human subject. Scientific reports. 15(1):29292. eng.

Chaurasia RK, Shirsath K, Mungse U, Bhat N, Khan A, Sapra B. 2025. FISH unveils a unified method for multi-marker biodose assessment. Scientific reports. 15(1):16994.

Chaurasia RK, Shirsath KB, Desai UN, Bhat NN, Sapra BK. 2022. Establishment of in vitro Calibration Curve for (60)Co-γ-rays Induced Phospho-53BP1 Foci, Rapid Biodosimetry and Initial Triage, and Comparative Evaluations With γH2AX and Cytogenetic Assays. Frontiers in public health. 10:845200. eng.

Constanzo J, Faget J, Ursino C, Badie C, Pouget J-P. 2021. Radiation-induced immunity and toxicities: the versatility of the cGAS-STING pathway. Frontiers in immunology. 12:680503.

Dagur PK, McCoy JP, Jr. 2015. Collection, Storage, and Preparation of Human Blood Cells. Current protocols in cytometry. 73:5.1.1-5.1.16. eng.

Delafont B, Carroll K, Vilain C, Pham E. 2018. Investigation of mixed model repeated measures analyses and non-linear random coefficient models in the context of long-term efficacy data. Pharmaceutical statistics. 17(5):515–526. eng.

Feinendegen LE. 2016. Quantification of Adaptive Protection Following Low-dose Irradiation. Health physics. 110(3):276–280. eng.

Feinendegen LE, Pollycove M, Neumann RD. 2007. Whole-body responses to low-level radiation exposure: new concepts in mammalian radiobiology. Experimental hematology. 35(4 Suppl 1):37–46. eng.

Feinendegen LE, Pollycove M, Neumann RD. 2009. Low-dose cancer risk modeling must recognize up-regulation of protection. Dose-response : a publication of International Hormesis Society. 8(2):227–252. eng.

Feinendegen LE, Pollycove M, Sondhaus CA. 2004. Responses to low doses of ionizing radiation in biological systems. Nonlinearity in biology, toxicology, medicine. 2(3):143–171. eng.

Fornalski KW, Adamowski Ł, Bugała E, Jarmakiewicz R, Krasowska J, Piotrowski Ł. 2024. Radiation adaptive response: the biophysical phenomenon and its theoretical description. Radiat Prot Dosimetry. 200(16–18):1585–1589. eng.

Guéguen Y, Bontemps A, Ebrahimian TG. 2019. Adaptive responses to low doses of radiation or chemicals: their cellular and molecular mechanisms. Cellular and molecular life sciences : CMLS. 76(7):1255–1273. eng.

He S, Lee W. 2022. Generalized linear mixed-effects models for studies using different sets of stimuli across conditions [Methods]. Frontiers in Psychology. Volume 13 - 2022. English.

Horn S, Barnard S, Rothkamm K. 2011. Gamma-H2AX-based dose estimation for whole and partial body radiation exposure. PloS one. 6(9):e25113. eng.

Jeggo P, Löbrich M. 2006. Radiation-induced DNA damage responses. Radiat Prot Dosimetry. 122(1–4):124–127. eng.

Kanani A, Krasowska J, Fornalski KW, Bevelacqua JJ, Welsh J, Mortazavi S. 2025. Adaptive Response: A Scoping Review of Its Implications in Medicine, Space Exploration, and Beyond. Dose-response : a publication of International Hormesis Society. 23(3):15593258251360051. eng.

Krasowska J, Fornalski KW. 2025. Can Adaptive Response Be Considered in Radiation Risk Assessment? Dose-response : a publication of International Hormesis Society. 23(2):15593258251341601. eng.

Lewis L, Crawford GE, Furey TS, Rusyn I. 2017. Genetic and epigenetic determinants of inter-individual variability in responses to toxicants. Current opinion in toxicology. 6:50–59. eng.

Liu Y, Hau KT, Liu H. 2024. Linear Mixed-Effects Models for Dependent Data: Power and Accuracy in Parameter Estimation. Multivariate behavioral research. 59(5):978–994. eng.

Lkhagvasuren E. 2017. Janeway’s immunobiology. Central Asian Journal of Medical Sciences. 3(1):100–101.

Mortazavi SM, Cameron JR, Niroomand-rad A. 2003. Adaptive response studies may help choose astronauts for long-term space travel. Advances in space research : the official journal of the Committee on Space Research (COSPAR). 31(6):1543–1551. eng.

Olivieri G, Bodycote J, Wolff S. 1984. Adaptive response of human lymphocytes to low concentrations of radioactive thymidine. Science (New York, NY). 223(4636):594–597. eng.

Pearson DD, Provencher L, Brownlee PM, Goodarzi AA. 2021. Modern sources of environmental ionizing radiation exposure and associated health consequences. Genome stability. Elsevier; p. 603–619.

Rogakou EP, Pilch DR, Orr AH, Ivanova VS, Bonner WM. 1998. DNA double-stranded breaks induce histone H2AX phosphorylation on serine 139. The Journal of biological chemistry. 273(10):5858–5868. eng.

Sankaranarayanan K, von Duyn A, Loos MJ, Natarajan AT. 1989. Adaptive response of human lymphocytes to low-level radiation from radioisotopes or X-rays. Mutation research. 211(1):7–12. eng.

Radioadaptive response and genomic instability: a phenotypic dichotomy of genome– environment interaction. International Congress Series; 2003: Elsevier.

Scott BR. 2014. Radiation-hormesis phenotypes, the related mechanisms and implications for disease prevention and therapy. Journal of cell communication and signaling. 8(4):341–352.

Shadley J, Wiencke J. 1989. Induction of the adaptive response by X-rays is dependent on radiation intensity. International journal of radiation biology. 56(1):107–118.

Sokolov MV, Smilenov LB, Hall EJ, Panyutin IG, Bonner WM, Sedelnikova OA. 2005. Ionizing radiation induces DNA double-strand breaks in bystander primary human fibroblasts. Oncogene. 24(49):7257–7265. eng.

Szumiel I. 1998. Monitoring and signaling of radiation-induced damage in mammalian cells. Radiat Res. 150(5 Suppl):S92–101. eng.

Tang FR, Loke WK. 2015. Molecular mechanisms of low dose ionizing radiation-induced hormesis, adaptive responses, radioresistance, bystander effects, and genomic instability. International journal of radiation biology. 91(1):13–27. eng.

Tapio S, Jacob V. 2007. Radioadaptive response revisited. Radiation and environmental biophysics. 46(1):1–12. eng.

Tigner A, Ibrahim SA, Murray I. 2020. Histology, white blood cell.

Tubiana M, Feinendegen LE, Yang C, Kaminski JM. 2009. The linear no-threshold relationship is inconsistent with radiation biologic and experimental data. Radiology. 251(1):13–22. eng.

Vaiserman A, Koliada A, Zabuga O, Socol Y. 2018. Health impacts of low-dose ionizing radiation: current scientific debates and regulatory issues. Dose-Response. 16(3):1559325818796331.

Wolff S. 1998. The adaptive response in radiobiology: evolving insights and implications. Environmental health perspectives. 106 Suppl 1(Suppl 1):277–283. eng.

